# RNA-seq highlights parallel and contrasting patterns in the evolution of the nuclear genome of holo-mycoheterotrophic plants

**DOI:** 10.1101/183202

**Authors:** M.I. Schelkunov, A.A. Penin, M.D. Logacheva

**Affiliations:** Institute for Information Transmission Problems, Russian Academy of Sciences, Moscow, Russia; Lomonosov Moscow State University, A.N Belozersky Institute of Physico-Chemical Biology, Moscow, Russia; Lomonosov Moscow State University, Faculty of Biology, Moscow, Russia

## Abstract

• While photosynthesis is the most notable trait of plants, several lineages of plants (so-called holo-heterotrophs) have adapted to obtain organic compounds from other sources. The switch to heterotrophy leads to profound changes at the morphological, physiological and genomic levels.

• Here, we characterize the transcriptomes of three species representing two lineages of mycoheterotrophic plants: orchids (*Epipogium aphyllum* and *Epipogium roseum*) and Ericaceae (*Hypopitys monotropa*). Comparative analysis is used to highlight the parallelism between distantly related holo-heterotrophic plants.

• In both lineages, we observed genome-wide elimination of nuclear genes that encode proteins related to photosynthesis, while systems associated with protein import to plastids as well as plastid transcription and translation remain active. Genes encoding components of plastid ribosomes that have been lost from the plastid genomes have not been transferred to the nuclear genomes; instead, some of the encoded proteins have been substituted by homologs. The nuclear genes of both *Epipogium* species accumulated mutations twice as rapidly as their photosynthetic relatives; in contrast, no increase in the substitution rate was observed in *H.monotropa*.

• Holo-heterotrophy leads to profound changes in nuclear gene content. The observed increase in the rate of nucleotide substitutions is lineage specific, rather than a universal phenomenon among non-photosynthetic plants.

## Introduction

The capability for photosynthesis is the iconic trait of plants and is of the highest importance to the biosphere. However, some plants, including several thousands of flowering plant species, obtain organic substances from sources other than photosynthesis (Merckx *et al.*, 2009; Westwood *et al.*, 2010). These plants acquire organic compounds either from associated fungi (myco-heterotrophy) or by parasitizing other plants. Most of these species combine photosynthesis and heterotrophy, but several hundred species have totally lost photosynthetic ability and become completely heterotrophic (holo-heterotrophs). The acquisition of heterotrophic ability has occurred in the evolutionary history of plants more than 50 times (Merckx *et al.*, 2009; Westwood *et al.*, 2010). The switch to heterotrophy leads to profound changes at the phenotypic level (reduction of leaves, loss of green colour, reduction of the vegetation period) that are highly parallel in different lineages. The genotypic alterations that underlie these changes are for the most part unclear. The difficulty of cultivating heterotrophic plants under experimental conditions hampers classic genetic and physiological studies. Advances in DNA sequencing permit the application of a genomic approach to elucidate the genetic changes associated with heterotrophy.

Genetic and genomic studies of heterotrophic plants are currently focused on two aspects. The first is the interaction of parasitic plants with their hosts and their adaptations to parasitism (e.g., Yang *et al.*, 2015). Extensive exchange of transcripts occurs between hosts and parasites (Kim *et al.*, 2014). On an evolutionary scale, a large number of horizontal gene transfer (HGT) events from hosts to parasites have been found in the organellar and nuclear genomes of parasitic plants (Bellot *et al.*, 2016; Li *et al.*, 2013). A recent large-scale survey of HGT in Orobanchaceae showed that the number of these events correlates positively with the degree of heterotrophy (Yang *et al.*, 2016). The second aspect is the evolution of organellar (mainly plastid) genomes. As expected, the plastomes of heterotrophic plants are reduced in size due to the loss of genes related to photosynthesis, and may retain only ~ 7.5% of the length of a typical plastome (Bellot & Renner, 2015). Despite the high degree of reduction, the plastomes of non-photosynthetic plants retain genes whose products are involved in translation, specifically transfer RNAs, components of the plastid ribosome and two other genes, *accD* and *clpP* (e.g., Wicke *et al.*, 2013; Barrett *et al.*, 2014; Schelkunov *et al.*, 2015; Bellot & Renner, 2015; Lam *et al.*, 2016). Although there are several considerations regarding the retention of the plastome, complete loss of the plastome is also apparently possible. At present, there are two known cases of such loss, one in *Polytomella*, a genus of unicellular algae (Smith & Lee, 2014), and one in the parasitic angiosperm *Rafflesia lagascae* (Molina *et al.*, 2014), although alternative explanations are possible in the latter case. Much less information on the mitochondrial genomes of non-photosynthetic plants is available, although there are indications that these genomes are not as extensively reduced in size (Fan *et al.*, 2016).

The changes in the nuclear genomes of non-photosynthetic plants have not been well studied. To date, the only work to deeply analyse the nuclear genomes of holo-heterotrophic plants was performed in Orobanchaceae, where a holo-heterotrophic species, *Orobanche aegyptiaca*, was compared with two of its relatives, one hemi-autotroph with obligatory parasitism and one hemi-autotroph with facultative parasitism (Wickett *et al.*, 2011). Surprisingly, the authors found evidence for conservation of the pathways responsible for chlorophyll synthesis. Additionally, in one study of the *Hypopitys monotropa* plastome, transcriptome analysis showed that many genes related to photosynthesis have been lost (Ravin *et al.*, 2016).

To obtain a more detailed understanding of the evolution of holo-heterotrophic plants, in this work, we analyse the nuclear genomes of *Epipogium aphyllum* and *E.roseum* (Orchidaceae) and *H.monotropa* (Ericaceae) using transcriptome sequencing. These plants are good models for studying the characteristics of holo-heterotrophic plants since their plastomes are among the most reduced in size (Schelkunov *et al.*, 2015; Logacheva *et al.*, 2016) (19-35 Kb versus approximately 150 Kb in typical photosynthetic plants). Therefore, we expect that the nuclear genomes of these species may also differ profoundly from the genomes of photosynthetic plants and will therefore allow us to highlight characteristics that are specific to the genomes of holo-heterotrophic species.

## Materials and methods

### Sample collection, library preparation and sequencing

Information on the specimens used for transcriptome sequencing was reported earlier by Schelkunov et al. 2015 for *E.aphyllum* (“White Sea” sample) and *E.roseum* (“Vietnam 2” sample) and by Logacheva *et al.* 2016 for *H.monotropa.* RNA was extracted using the Qiagen RNeasy Plant Mini kit (Qiagen, Netherlands). To allow the characterization of non-polyadenylated transcripts (e.g., plastid and mitochondrial), we prepared libraries from ribosomal RNA-depleted samples using the Ribo-Zero plant leaf kit (Illumina, USA). Detailed information on the libraries and sequencing settings, as well as links to on-line databases where the reads and assembled sequences are deposited, is provided in Table S1.

### Choice of datasets of photosynthetic plants

To compare the holo-heterotrophic species with typical photosynthetic plants, we obtained data from several RNA-seq experiments and from the genome assemblies of the plants that were available as of 2015. Although many transcriptomes have been sequenced, the corresponding data exhibit certain shortcomings, such as a low sequencing depth, the use of technologies that introduce errors in homopolymer regions, or the sequencing of organs (e.g., roots) in which the set of expressed genes may differ from that in above-ground organs. In view of these considerations, we chose six photosynthetic species, including three for comparison with *Epipogium* and three for comparison with *H.monotropa*. The names and sources of the datasets on photosynethetic plants are listed in Tables S1 and S2. In the case of RNA-seq data, we assembled the transcriptomes using exactly the same methods that were used for the transcriptomes of *Epipogium* and *Hypopitys*.

### Transcriptome assembly and postprocessing

Reads were trimmed with Trimmomatic 0.32 (Bolger *et al.*, 2014). Trimming involved the removal of sequencing adapters, bases with a Phred quality of less than 3 from both the 5’ and 3’ ends of reads, and bases from the 5’ ends of reads starting from a region with 5 consecutive bases with an average score of less than 10 (trimming with a sliding window). Additionally, reads that had average quality of less than 20 after trimming and reads that after trimming became shorter than 30 bp were removed. Assembly was performed using Trinity version r20140717 (Grabherr *et al.*, 2011). Digital normalization of coverage to 50× was switched on. After assembly, we performed filtration, removing minor isoforms, low-coverage transcripts and contamination. We defined major isoforms as the isoforms to which the highest number of reads (estimated using RSEM (Li & Dewey, 2011)) were mapped relative to other isoforms. After removing minor isoforms, we filtered low-coverage transcripts by mapping reads to contigs using CLC Assembly Cell 4.2 and retaining only transcripts with an average coverage of at least 3. Potential CDSs in the transcripts were then determined using TransDecoder version r20140704 (Haas *et al.*, 2013). The criteria for considering an ORF to be a potential CDS consisted not only of hexanucleotide frequencies, which are employed in TransDecoder by default, but also all ORFs that possessed domains from the Pfam-A or Pfam-B databases. The minimum CDS length was decreased from the default of 100 amino acids to 30 amino acids. When there were several potential CDSs from a transcript, only the longest one was taken. Then, to remove contamination, we performed BLASTP 2.2.29+ (Camacho *et al.*, 2009) alignment (maximum allowed evalue 10^−5^, word size 3, the low-complexity filter switched off) of the translated CDS sequences against the NCBI NR database and against proteins of related plants whose genomes had been sequenced. A CDS was considered to represent contamination if the BLASTP match with the highest bit score was to a species that was not from Streptophyta. Conversely, if the best match was to Streptophyta member or the sequence presented no BLASTP matches, the CDS was considered to belong to a plant. Statistical parameters of the assemblies, such as N50 values and the number of sequences longer than 1000 bp, were calculated using custom scripts after each filtering step described above. Estimation of the completeness of the assemblies was performed with CEGMA 2.5 (Parra *et al.*, 2007). To evaluate gene expression without contamination, we calculated FPKM values using RSEM for transcripts that contained CDSs that were determined not to have arisen from contamination. Minor isoforms of the transcripts were also used for the analysis; the FPKM values provided in Table S3 are the sums of all gene isoforms.

### Transcriptome annotation and Gene Ontology analysis

Three types of annotations of the CDSs were performed. The first was the computation of 1-1 orthologs from the CDSs of each species by reciprocal alignment with *Arabidopsis thaliana* proteins from TAIR10 database (Berardini *et al.*, 2015) using BLASTP (parameters are the same as indicated above). Also we performed GO annotation using results of BLASTP alignment of all proteins against the NCBI NR (parameters as above) and annotated them with B2G4Pipe 2.5, a command-line wrapper for Blast2GO (Conesa *et al.*, 2005). Then we assigned KEGG Orthology identifiers to the CDSs with the GhostKoala server (Kanehisa *et al.*, 2016b). The orthogroups in the studied species were calculated separately for Orchidaceae and Ericales using OrthoMCL 2.09 (Li *et al.*, 2003) (inflation parameter 1.5).

For GO term enrichment analysis, we used a set of custom scripts written in Perl and R. Utilizing the results of the GO annotation performed with Blast2GO and a graph of GO terms (http://purl.obolibrary.org/obo/go/go-basic.obo, last accessed 15 April 2015), the scripts provide all GO terms corresponding to a gene, including terms that are paternal (i.e., with higher hierarchical levels) to those provided by Blast2GO. Comparison of the numbers of genes with specific GO terms between a pair of species was then performed via a series of Fisher’s exact tests (one for each GO term). GO terms that were not assigned to any genes in either species were excluded from the analysis. After the Fisher’s tests were performed, the scripts performed group comparisons between holo-heterotrophic and photosynthetic plants for each GO term, by taking all p-values for each pairwise comparison between pairs of species within the group (Ericales or Orchidaceae). The scripts then conducted Bonferroni correction for multiple testing to evaluate the statistical significance of the differences between these groups separately for Orchidaceae and Ericales. Next, another round of correction for multiple hypothesis testing was performed, taking into account the fact that an individual Fisher’s test was performed for each GO term. This correction was performed via the method of Benjamini-Yekutieli (Yekutieli & Benjamini, 2001).

Since our goal was the analysis of nuclear genes, we excluded genes encoded in the plastid and mitochondrial genomes from the enrichment analysis by excluding all genes that were 1-1 orthologs of plastid and mitochondrial genes of *Arabidopsis thaliana.* Some plastid and mitochondrial transcripts may not have been present in the assemblies. To compensate for this, in Table S3, we indicated a plastid gene as present in the plastome of a studied species not only when it was found through the 1-1 ortholog method, but also when it was present in the annotation of the plastome of that species provided in GenBank (accessions are provided in Table S2). Because the plastome of *O. italica* is not available, we employed the plastome of *Habenaria pantlingiana*, the only species with a characterized plastome from subtribe Orchidiae, which includes *O. italica*.

### Analysis of substitution rates and selective pressure

To estimate the average selective pressure and substitution rates in the genomes of the studied species, we concatenated the sequences of all genes from orthogroups containing exactly one gene from each species. The concatenated sequences were then aligned using TranslatorX 1.1 (Abascal *et al.*, 2010). As a tool for the alignment of the amino acid sequences by TranslatorX, we used Muscle (Edgar, 2004). The topologies of the phylogenetic trees for Orchidaceae and Ericales were obtained from the APG III classification (THE ANGIOSPERM PHYLOGENY GROUP, 2009). dN/dS and substitution rate estimation was performed based on the alignment and the tree using PAML 4.8 (Yang, 2007), employing a branch model with free ratios, the Gy94_3×4 codon model and removal of all columns with at least one gap in the alignment.

To compare the magnitude of the selective pressure acting on individual genes in heterotrophic and autotrophic plants, the sequences of the genes were aligned using TranslatorX and Muscle as described above, and dN/dS ratios were calculated using PAML in the branch model. For the Orchidaceae, two calculations were performed. In the first calculation, one dN/dS was allowed for the branches of autotrophic plants; another dN/dS was allowed for the terminal branches of *E.aphyllum* and *E.roseum*; and a third dN/dS was allowed for the branch of their common ancestor. The second calculation was performed allowing one dN/dS for the common ancestor of *E.aphyllum* and *E.roseum* and a second dN/dS for all other branches, including autotrophic species, *E.aphyllum* and *E.roseum*. The P-values for the difference in dN/dS between the terminal branches of *Epipogium* and the branches of autotrophic species were calculated using the likelihood ratio test. Allowing an individual dN/dS for the branch of the common ancestor of *Epipogium* was beneficial, as photosynthetic ability was lost on that branch, and it is unclear whether to group it with autotrophic or heterotrophic species. Similar calculations were performed for Ericales, allowing different dN/dS ratios for the *H.monotropa* branch and other branches the first time and demanding a single dN/dS for all branches the second time. Since *H.monotropa* was the only heterotrophic species from Ericales employed in this study, its terminal branch partially included its autotrophic ancestor. Thus, the values of dN/dS provided for *H.monotropa* in Table S3 describe selective pressure partially before and partially after the loss of photosynthetic ability. dN/dS was calculated only for genes that were present in all 5 species of Orchidaceae or in all 4 species of Ericales.

### Analysis of protein targeting to organelles

To estimate the number of proteins targeted to various organelles, we predicted transit peptides using TargetP 1.1 (Emanuelsson *et al.*, 2000) and DualPred (Saravanan & Velan Lakshmi, 2015). Unlike TargetP, DualPred is also capable of predicting proteins that are dually targeted to mitochondria and plastids. TargetP classifies predictions into five “reliability classes”, where the most confident predictions belong to the first class, and the least confident predictions belong to the fifth class. We considered a protein to be potentially targeted to an organelle if its transit peptide exhibited a reliability class of four or less. For DualPred, we considered a protein to be dually targeted to plastids and mitochondria if it presented a dual-targeting score of at least 0.5, as suggested by the author of DualPred in a personal communication. Prediction of transit peptides was performed only for proteins whose genes exhibited a completely assembled 5’-end according to TransDecoder and were assigned at least one GO term.

To predict the targeting of ribosomal proteins, we utilized a more elaborate technique. It has been demonstrated that some proteins are dually targeted to plastids and mitochondria not because they have one transit peptide that allows them to enter both organelles, but because their mRNAs exhibit alternative translation start sites, resulting in proteins with different transit peptides (Mitschke *et al.*, 2009). To search for transit peptides that may originate from alternative translation, we truncated all of the ribosomal proteins under analysis before the first methionine occurring after the first 25 amino acids (as in Mitschke *et al.*, 2009) and performed TargetP and DualPred analyses for these shortened versions of the proteins as well. Additionally, all alternative isoforms of the ribosomal proteins were analysed. When DualPred suggested that a protein was dually targeted and TargetP suggested either plastid or mitochondrial targeting, we considered this to represent a non-contradictory prediction suggesting dual targeting.

The Pfam families to which the ribosomal proteins belonged were determined by aligning *Arabidopsis thaliana* proteins to all families from the Pfam database, version 31.0 (Finn *et al.*, 2016), using the HMMER server (Finn *et al.*, 2015). To identify proteins that belonged to these families in *E.aphyllum*, *E.roseum* and *H.monotropa*, we conducted a search with hidden Markov models of these families using the hmmscan tool from HMMER package, version 3.1b1 (Eddy, 1998). The search was performed among all of the CDSs from transcripts, not only the longest ones, to make detection in polycistronic mitochondrial mRNAs possible. Orthologous proteins in the photosynthetic species employed for comparison were determined as belonging to orthogroups that contained the proteins identified for *E.aphyllum*, *E.roseum* and *H.monotropa*.

### Graphic representation of results

Phylogenetic trees were drawn with TreeGraph 2.9.2 (Stöver & Müller, 2010). Maps of metabolic and signalling pathways were built the KEGG site (Kanehisa *et al.*, 2016a) (accessed 22 October 2015).

**Table 1.** Brief statistics of the transcriptome assemblies.

|  | *Epipogium aphyllum* |  | *Epipogium roseum* |  | *Hypopitys monotropa* |  |
| --- | --- | --- | --- | --- | --- | --- |
|  | All transcripts | CDSs* | All transcripts | CDSs* | All transcripts | CDSs* |
| Number of sequences | 992 338 | 20 958 | 1 336 170 | 19 026 | 217 166 | 13 276 |
| Number of sequences longer or equal to 1000 bp | 39 403 | 4 321 | 28 284 | 3 259 | 33 560 | 5 421 |
| Total length of sequences | 290 947 719 | 12 988 731 | 321 048 790 | 10 721 418 | 116 614 033 | 13 496 709 |
| N50 | 371 | 1 173 | 257 | 990 | 1 199 | 1470 |
| Median length of sequences | 178 | 306 | 164 | 315 | 237 | 807 |

\*Here, the CDS is defined as the longest ORF in the major isoforms of the transcripts that were assigned at least one GO term (low-coverage transcripts and contaminating transcripts are not considered).

## Results and Discussion

### Characteristics of transcriptome assembly

The statistics of transcriptome assembly are provided in Table 1 (for details, see Table S4). Simple statistical measures, such as N50 values and the mean contig length, were smaller than those in many studies involving transcriptome assemblies, mainly because we analysed contigs with a minimum length of 100 bp instead of 200 bp (the default cut-off in the Trinity assembler). Several genes of interest (for example, the plastid genes *rpl32* and *rpl36*) are shorter than 200 bp, and use of the default cutoff would incur a risk of missing their transcripts. The assembly statistics were also biased by contamination. The symbiosis of mycoheterotrophic plants with fungi may be quite deep; for example, the rhizome of *E.aphyllum* has been described as “heavily and permanently infected” (Rasmussen, 1995). Despite the fact that we sequenced RNA from the above-ground parts of the plants, we observed a large number of bacterial and fungal transcripts in the assemblies (Fig. S1), especially in *E.roseum*, in which they represented ~80% of contigs that we were able to taxonomically classify by BLAST. In *E.aphyllum* and *H.monotropa*, the corresponding values were approximately 10% and 5%, respectively. This difference may reflect the correlation between soil microbiome biomass and climate (*E.aphyllum* and *H.monotropa* were collected from colder regions than *E.roseum*). Among the transcriptomes of photosynthetic plants used for comparison, the transcriptome of *Orchis italica* was also highly contaminated. Sequences originating from fungi and bacteria were discarded prior to further analysis. Estimation of the completeness of the assemblies based on the set of genes that are expected to be present in all eukaryotic genomes (Parra *et al.*, 2007) showed that > 90% of these genes were at least partially assembled (Table S5). Notably, the genomes were assembled less completely than the transcriptomes on average, with a median of 95.6% of the genes being assembled at least partially in the transcriptome assemblies, compared with 90.8% of the genes in the genome assemblies. These results show that, given sufficient coverage, RNA-seq is as good as complete genome sequencing in terms of the number of retrieved genes, while being less costly and using an assembly process that is computationally faster.

### Gene retention and reduction

Plastid-targeted proteins carry specific amino acid sequences known as targeting signals or transit peptides that interact with the translocon system and enable the import of these proteins into plastids. Thus, we expected that in non-photosynthetic plants, the fraction of proteins with plastid-targeting signals relative to other proteins will be decreased. However, a comparison revealed a more complex situation. In *Epipogium*, the fraction of proteins targeted to plastids relative to the total number of proteins is approximately 2 times lower than in autotrophic orchids on average, whereas the fraction of proteins that are targeted to mitochondria or the endoplasmic reticulum is approximately the same. In contrast, in *H.monotropa*, the fraction of proteins targeted to plastids does not appear to be decreased. However, because the fraction of plastid-targeted proteins differs greatly between the two photosynthetic Ericales species that we used for comparison (Table S6), these results should be treated with caution, as they may be biased by lineage-specific genome duplications and/or by differences in the quality of the assemblies. To obtain a deeper understanding of the patterns of gene reduction, we performed Gene Ontology (GO) enrichment analyses. GO analysis of *Epipogium aphyllum* and *E.roseum* versus three photosynthetic orchids revealed that the genes associated with 60 GO terms in *Epipogium* were underrepresented, while the genes associated with 38 terms were overrepresented, with q-values ≤ 0.05. All of the overrepresented GO terms are related to genes associated with mobile elements. This result is presumably caused by methodological differences, as the *Epipogium* transcriptomes were sequenced without selection of polyadenylated transcripts, whereas the transcriptomes of *Cymbidium ensifolium* and *O. italica* represented polyA fractions. Thus, mobile elements, whose RNAs are not polyadenylated (Chang & Schulman, 2008) are expected to be overrepresented in *Epipogium*. In the genome of *Phalaenopsis equestris*, mobile elements are masked as repeats, producing a similar effect. Among the underrepresented GO categories, almost all were related to photosynthesis and plastids. The least underrepresented GO terms were general and difficult to interpret (e.g., “Single-organism metabolic process” and “Membrane”). The most underrepresented GO terms are listed in Table 2; for a complete list, see Table S3.

**Table 2.**
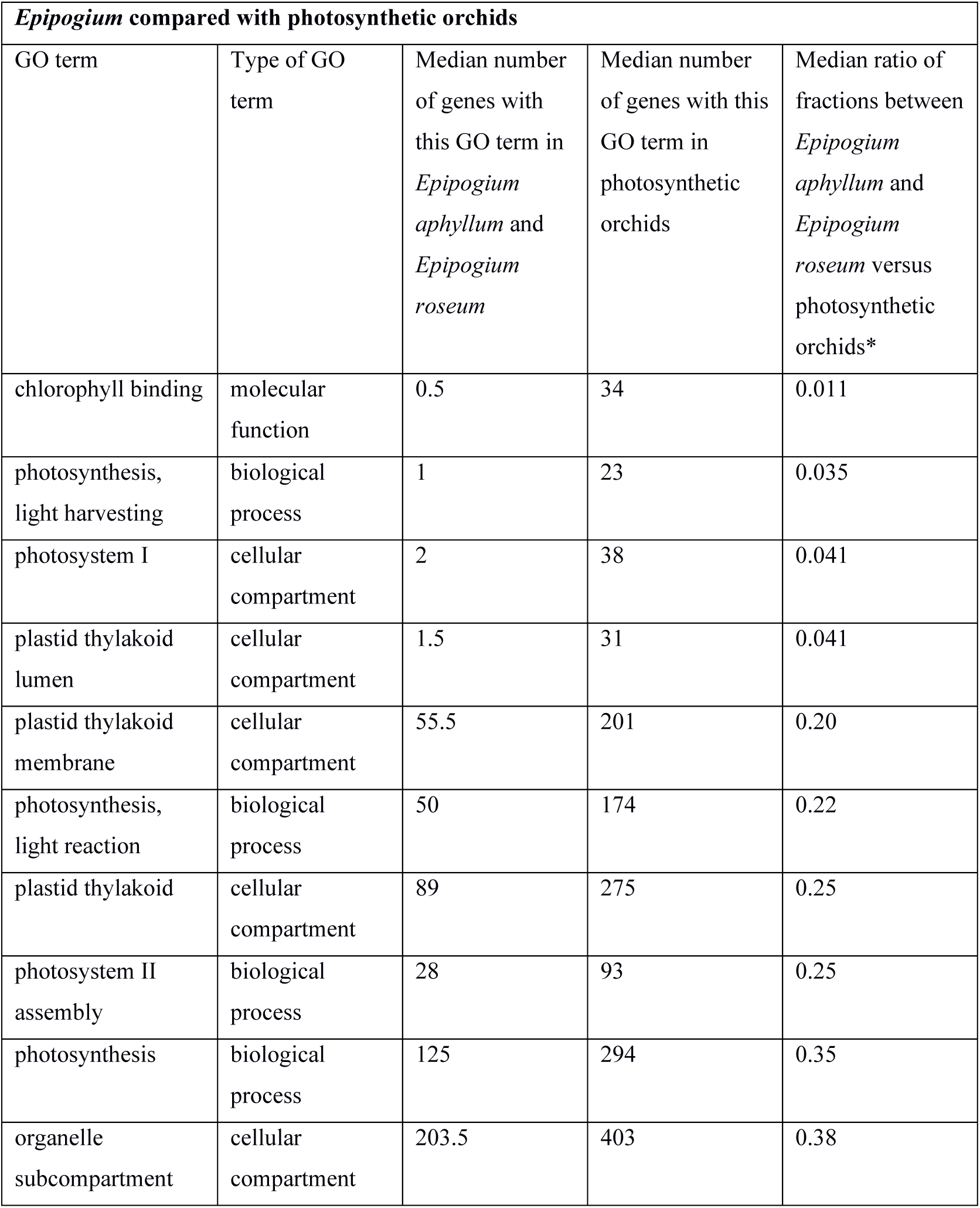
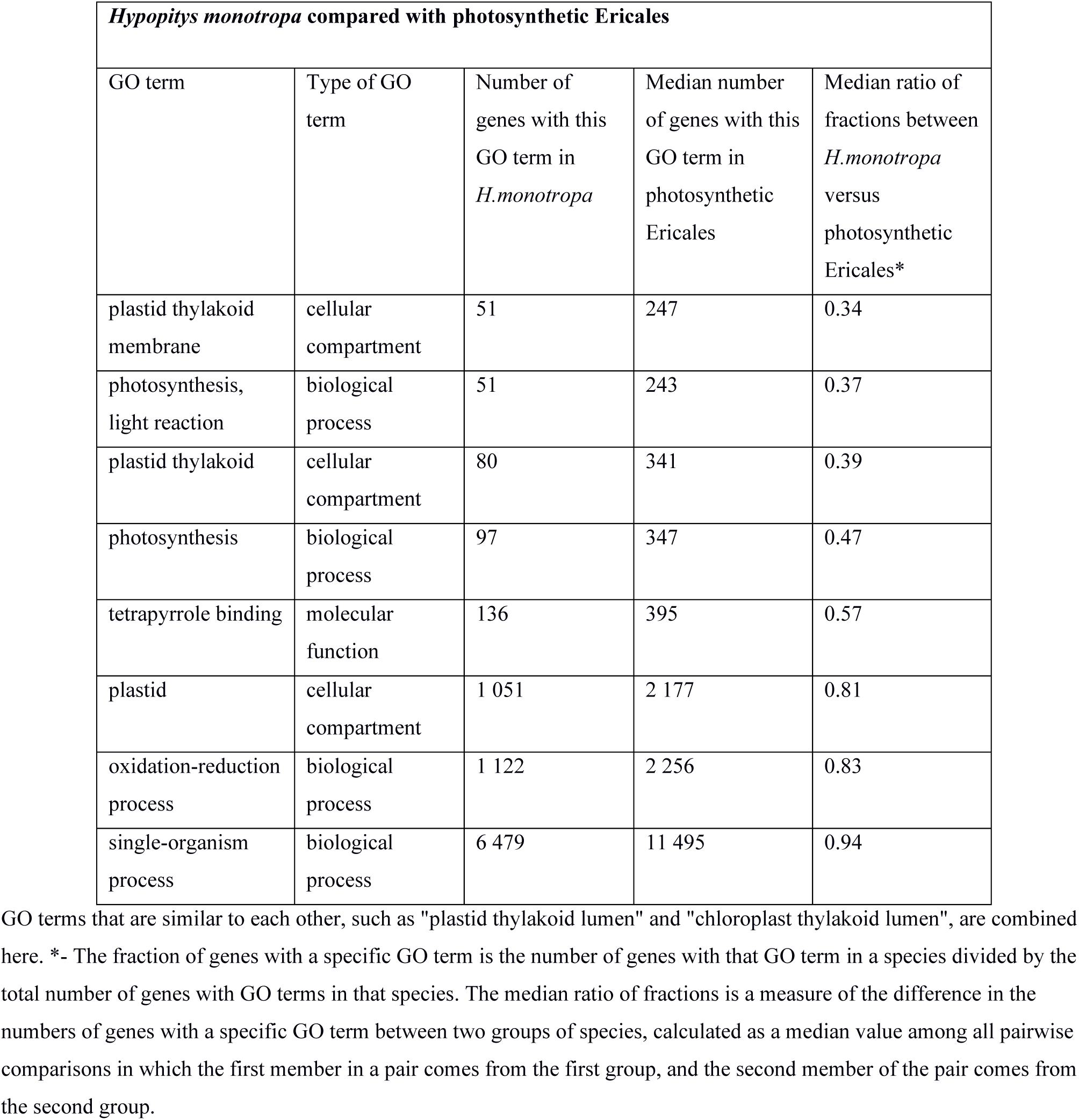
GO terms showing the greatest differences in the fraction of genes between mycoheterotrophic species and their photosynthetic relatives.

GO enrichment analyses between *H.monotropa* and three photosynthetic Ericales showed similar results; 17 GO terms were overrepresented, and 9 are underrepresented. The overrepresented terms were related to mobile elements, and the underrepresented terms were related to photosynthesis and plastids (Table 2; Table S3).

Notably, we did not observe changes in the list of GO terms other than related to photosynthesis and plastids, despite dramatic differences in the morphology and physiology of the studied species. This finding indicates that these differences are controlled not at the level of the presence or absence of specific genes, but rather by the regulation of gene expression. To gain insight into this regulation, more detailed transcriptome data are required. Alternatively, these morphological and physiological changes may have originated from the disruption or loss of a small number of genes which do not produce statistically significant results in GO enrichment analysis.

The statistical analysis of GO terms is quite rough method and may not reflect differences at the level of individual genes. Thus, we also searched for orthologs of genes that are known to participate in processes occurring in the plastids in the model plant *Arabidopsis thaliana*. As expected, genes related to photosynthesis have been lost from nuclear genomes of *Epipogium* and *H.monotropa* (Table 3). In particular, no nuclear-encoded components of the plastid electron transfer chain (*PSB* and *PSA* genes) were found, which is consistent with the absence of plastid-encoded *psa* and *psb* genes from the plastomes of all three species. Components of the light-harvesting antenna (*LHC* genes) were completely absent from *H.monotropa*; one such gene (*LHCA4*) was present in the *E.aphyllum* transcriptome, but its expression (measured in FPKM) was ~7-25-fold lower than in photosynthetic orchids. Nevertheless, this gene exhibited a full-length open reading frame that has evolved under negative selection (dN/dS 0.11). Since the product of this gene participates in chlorophyll binding, its retention may be in some way related to the retention of chlorophyll synthesis in *Epipogium*, as described below. Additionally, as shown in Table S7, a few transcripts of genes that encode proteins in the plastid electron transfer chain were found in the transcriptomes of *Epipogium aphyllum*, *E.roseum* and *H.monotropa*. All of the observed sequences were shorter than 50% of the length of their orthologs in photosynthetic plants and are likely to be pseudogenes. The apparent partial conservation of enzymes in the Calvin cycle is due to that some of these enzymes are involved in metabolic processes that are not related to photosynthesis, e.g., glycolysis. Many genes that encode Calvin cycle enzymes belong to small gene families (e.g., *GAPDH*, *PGK*, *TKL*, *TPI*) in which different members encode proteins that exhibit the same enzymatic activity (isoenzymes) but show different cellular localizations and act in different pathways. For example, the GAPDH and TPI proteins function in both glycolysis (cytosolic isoenzymes) and the Calvin cycle (plastidic isoenzymes). We assume that most of the transcripts corresponded to cytosolic isoenzymes. Consistent with this assumption, 8 of 10 of the transcripts found in *Epipogium* and 7 of the 14 transcripts found in *H.monotropa* did not have plastid-targeting signals (Table S7), and those that have (*TKL*, *RPI*) exhibit additional functions in plastids that are not related to the Calvin cycle. Expression of genes encoding proteins that act exclusively in photosynthesis (e.g., rbcS, SBPase) was absent. The nuclear-encoded components of plastid ATP synthase have been completely lost, as well as the plastid-encoded components. The sigma subunits of plastid-encoded RNA polymerase (PEP) and PEP-associated proteins have also been lost. PEP is involved in the transcription of genes related to photosynthesis, unlike nuclear-encoded RNA polymerase (NEP), which mainly transcribes plastid genes that are unrelated to photosynthesis (Shiina *et al.*, 2005). The redox-sensitive components of the plastid translocon, which is the system responsible for the import of proteins from the cytoplasm to plastids, have also been lost, with the exception of TIC32 of *Hypopitys*.

**Table 3.** Number of selected photosynthesis-related nuclear genes and nuclear genes encoding proteins involved in plastid functions other than photosynthesis in *Epipogium aphyllum* and photosynthetic orchids as well as *Hypopitys monotropa* and photosynthetic Ericales.

| Genes | Number in *E.aphyllum* | Median number in photosynthetic orchids | Number in *H.monotropa* | Median number in photosynthetic Ericales |
| --- | --- | --- | --- | --- |
| Photosynthesis-related |  |  |  |  |
| Components of photosystem I | 0 | 9 | 0 | 8 |
| Components of photosystem II | 0 | 9 | 0 | 8 |
| Components of electron transfer chain (others than PSI and PSII) | 0 | 5 | 0 | 6 |
| Light-harvesting complex | 1 | 10 | 0 | 8 |
| Calvin cycle | 10 | 20 | 14 | 21 |
| Sigma subunits of PEP and PEP-associated proteins | 0 | 14 | 1 | 14 |
| Plastid ATP synthase | 0 | 3 | 0 | 3 |
| Non-photosynthesis-related |  |  |  |  |
| Plastid ribosome | 33 | 34 | 31 | 31 |
| Clp subunits | 11 | 12 | 10 | 10 |
| ACC subunits | 5 | 6 | 4 | 7 |
| Plastid translocon | 15 | 20 | 11 | 18 |

In contrast to genes encoding proteins that are necessary for photosynthesis, genes that are responsible for other functions of plastids have been retained (Table 3). In particular, components of clp-protease and acetyl-CoA carboxylase whose counterparts (clpP and accD) are encoded in the plastome have been retained. Consistent with this finding, NEP, which transcribes genes not related to photosynthesis, has been retained in both species, as have most components of the plastid ribosome (but see the discussion below). We also found transcripts of most proteins responsible for the replication and repair of the plastome. However, this situation is more complex because many of these proteins are targeted not only to plastids but also to mitochondria and the nucleus, and information on these proteins is sometimes inconsistent between different experiments (Tanz *et al.*, 2013; Huang *et al.*, 2013).

As shown above, the genomes of *Epipogium* and *H.monotropa* encode a number of proteins that must be imported into plastids. Accordingly, the components of the plastid translocons for both the outer- and inner-envelope membranes must be retained. Recent studies in *A. thaliana* have shown that the plastid-encoded protein ycf1 (TIC214) plays an essential role in plastid translocation and that it acts at the inner membrane (Kikuchi *et al.*, 2013). However, the *ycf1* gene is absent from both the *Epipogium* and *H.monotropa* plastomes. A transcript similar to *ycf1* was only found in the transcriptome assembly of *E.aphyllum*, in which it carried a signal for targeting of the protein to mitochondria. Homologs of TIC100 and TIC56, which encode proteins that interact with Ycf1 within the TIC complex, were also absent from all three species. It should be noted that Ycf1 and its interacting proteins are also absent from several photosynthetic species, including grasses and *Vaccinium macrocarpon*, which raises a question regarding the universality of the function of Ycf1 (de Vries *et al.*, 2015). The current model of TIC postulates the existence of two complexes. The first, referred to as the “photosynthetic-type” complex, consists of Ycf1, TIC56, TIC100 and TIC20-I and is a major TIC complex that functions in most land plants to import proteins involved in photosynthesis. The second complex is a “non-photosynthetic-type” complex, which imports proteins that are not related to photosynthesis (Nakai, 2015b). It has been hypothesized that the switch from the major to the alternative system of protein import occurred in grasses (Nakai, 2015b; Nakai, 2015a) and that the major system then degraded. The pattern of gene loss and retention observed in *Epipogium* and *H.monotropa* suggests that this could also be the case in these species. The existence of two complexes, one of which mainly imports photosynthetic proteins, while the other is non-photosynthetic, has also been postulated for the outer translocon TOC (Nakai, 2015b; Nakai, 2015a). *Epipogium* and *H.monotropa* possess orthologs of TOC proteins, but not a complete set (Table 2), suggesting that only one of these complexes is retained in these plants.

Annotation of the transcriptomes using the KEGG database of biological pathways revealed reductions in the number of genes associated with several other pathways, some of which are common to *Epipogium* and *H.monotropa*, while others are lineage-specific (Figs. S2-S5). For example, a reduction in the number of proteins involved in light reception and circadian rhythms was observed; although this reduction was found in *Epipogium* as well as in *Hypopitys*, it was notably different in these holo-heterotrophic plants (Fig. S2). In particular, *Epipogium* appeares to lack genes encoding the photoreceptors phytochrome B and cryptochrome, whereas *H.monotropa* lacked several proteins (LHY, ZTL and GI) that regulate the circadian clock. The absence of these proteins may be related to the distinctive lifestyles of these plants, which spend most of their life cycle completely underground and appear above-ground only for several weeks during blossoming (Bjorkman, 1960; Yagame *et al.*, 2007; Taylor & Roberts, 2011), and may therefore have different requirements for the regulation of circadian rhythms than typical autotrophic plants. However, both *Epipogium* and *H.monotropa* have retained essential elements of circadian clock regulation (HY5, ELF3), photoreceptor phytochrome A and proteins interacting with them (PIF, COP1), indicating that the core proteins that regulate plant development under the influence of light have been conserved. In addition, we found a reduction in the number of genes associated with the carotenoid (Fig. S3) and phylloquinone (Fig. S4) biosynthetic pathways. As carotenoids are a component of light-harvesting antennae, the reduction in the number of genes related to carotenoid biosynthesis is presumably linked to the disappearance of photosynthesis. One interesting and unexplained observation was a reduction in the thiamine synthesis pathway that affected *Epipogium*, but not *H.monotropa* (Fig. S5). Another contrasting characteristic was the retention of the chlorophyll *a* synthesis pathway in *Epipogium*, whereas only the gene responsible for the first step in this pathway has been retained in *H.monotropa* (Fig. 1). In *H.monotropa*, the dN/dS for the only retained gene in this pathway was significantly higher than in autotrophic Ericales (p-value of 0.02 by the likelihood ratio test), showing a value of 0.27 in *H.monotropa* versus 0.11 in its autotrophic relatives. Among the 5 considered genes in *Epipogium*, only 1 showed increased dN/dS with a significant p-value (< 0.05 by the likelihood ratio test with Bonferroni correction). The mean dN/dS for these 5 genes in *Epipogium* was only slightly higher than the mean in its photosynthetic relatives, at 0.11 versus 0.05, respectively (Table S7). However, all of the genes in this pathway were expressed at levels many times lower than in photosynthetic species (Table S7), suggesting that it could be not functional being the remnant of one that was active in a photosynthetic ancestor of *Epipogium*. Alternatively, chlorophyll *a* could indeed be synthesized in these species, where it may act in cellular processes that are unrelated to photosynthesis. Chlorophyll *a* has been found in many heterotrophic plants via chromatography (Cummings & Welschmeyer, 1998) (notably, among these species is *Monotropa uniflora*, a close relative of *H.monotropa*); transcriptome sequencing in parasitic *Orobanche aegyptiaca* has also demonstrated the presence of genes responsible for chlorophyll *a* synthesis, with no signs of relaxed selection (Wickett *et al.*, 2011). It has been shown that in *A. thaliana*, pheophorbide *a*, a product of chlorophyll *a* catabolism, causes cell death in a light-independent manner (Tanaka & Tanaka, 2006; Hirashima *et al.*, 2009). Conservation of this mechanism in *Arabidopsis* and non-photosynthetic plants could explain the conservation of the chlorophyll *a* synthesis pathway.

**Figure 1.**
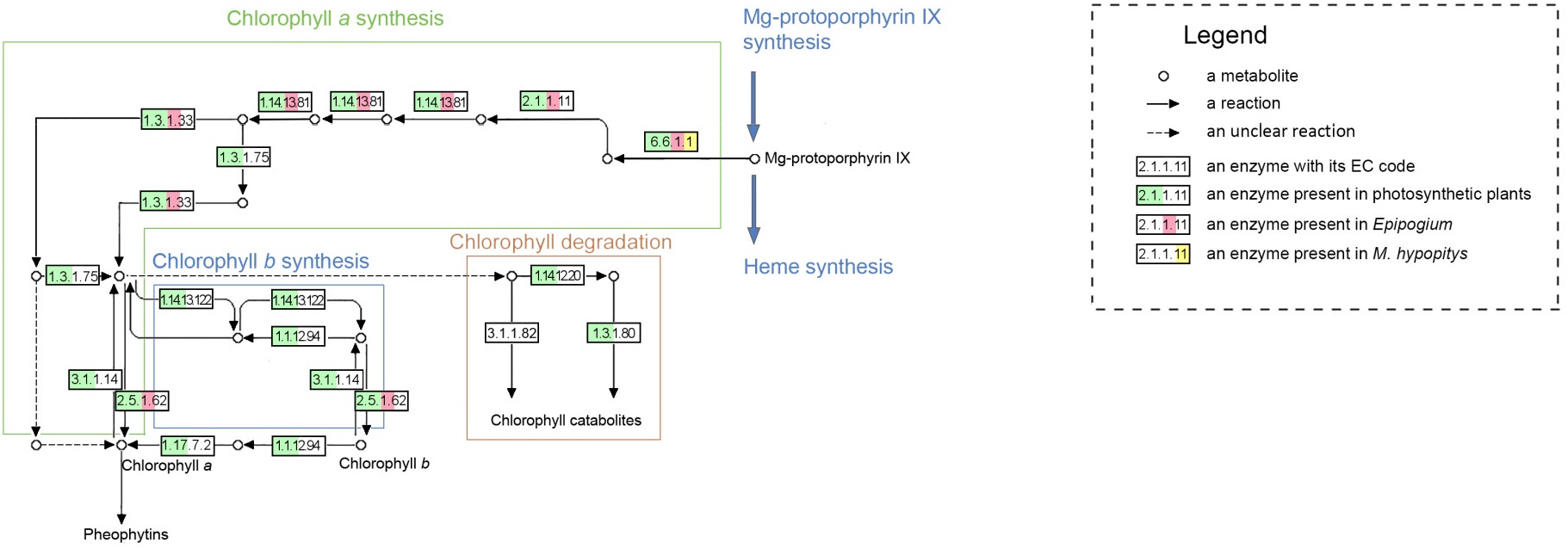
Map of the chlorophyll metabolism pathway showing proteins present in *Epipogium*, *Hypopitys monotropa* and green plants. A protein is depicted as present in a specific plant group if its transcript was observed in a transcriptome of at least one species of that group.

Despite the differences discussed above, *Epipogium* and *H.monotropa* show striking parallelism in the reduction of their nuclear genes, considering that these plants lost autotrophic ability independently and that they diverged approximately 150 million years ago (Hedges *et al.*, 2015).

Quantitative estimates of this parallelism can be biased by differences in assembly quality and by lineage-specific gene losses that are not related to heterotrophy. To obtain an unbiased estimate, we considered only the genes that presented one-to-one orthologs in the *Arabidopsis* genome and were found in all 6 photosynthetic species used for comparison (4888 genes in total). Among these genes, 818 were absent in *E.aphyllum*, and 745 were absent in H.monotropa. In *E.roseum*, 996 genes were absent, which presumably reflects the lower quality of the assembly obtained for this species. For *E.aphyllum* and *E.roseum*, the overlap between the lost genes was 67%, whereas for *E.aphyllum* and *H.monotropa*, it was 49%, and for *E.roseum* and *H.monotropa*, it was 45%. If the losses occurred randomly, we would expect a much lower (~16%) percentage of common losses. According to the hypergeometric test, the p-value for the null-hypothesis that losses are uncorrelated was <10^−100^ for both the *E.aphyllum*/*H.monotropa* and *E.roseum*/*H.monotropa* pairs.

### Mutation accumulation rates

Variation in mutation accumulation rates is a phenomenon that is widely observed in plants (e.g., Bromham *et al.*, 2015). In particular, it has been found that in parasitic plants, the mutation accumulation rate is increased not only in plastome, as expected for heterotrophic plants, but also in nuclear and mitochondrial genomes (Bromham *et al.*, 2013). Concerning mycoheterotrophic plants, information on mutation rates is mostly limited to plastomes, which show increased substitution rates with only rare exceptions (Logacheva *et al.*, 2014; Barrett *et al.*, 2014). Our previous studies of *Epipogium* and *H.monotropa* indicated that the mutation accumulation rate in plastid genes, at both synonymous and in nonsynonymous sites, was increased in comparison to the holo-autotrophic relatives of these species, but at different levels (approximately 20 and 2.5 times respectively) (Schelkunov *et al.*, 2015; Logacheva *et al.*, 2016). Characterization of transcriptome sequences allowed us to test whether this increase was confined to plastid genes. To calculate the mutation accumulation rate in the nuclear genomes, we used concatenated sequences of genes from orthogroups containing exactly one gene in each species. There were 4479 and 2547 of these orthogroups in Orchidaceae and Ericales, respectively. The mutation accumulation rates in both *Epipogium* species were approximately two times higher than the mutation rates in their photosynthetic relatives. The rates of nonsynonymous and synonymous mutation accumulation in *Epipogium* were increased proportionally (Figs. S6-S8). In contrast, the mutation accumulation rate in *H.monotropa* was not increased (Fig. 2).

**Figure 2.**
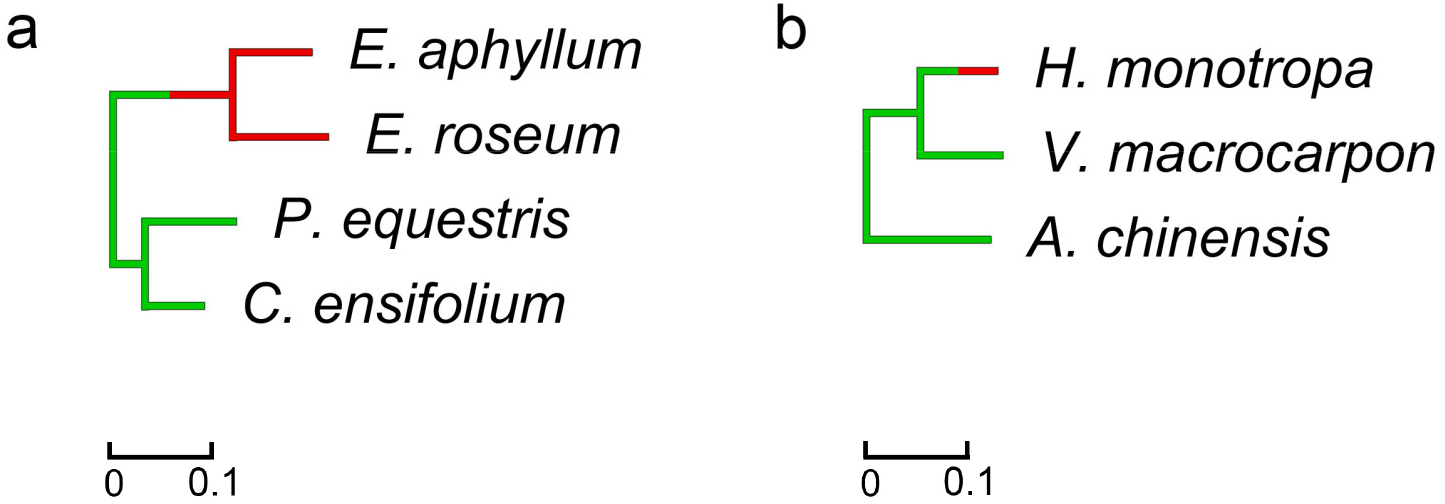
Mutation accumulation rates in nuclear genes of *Epipogium* (a) and *Hypopitys monotropa* (b). The branch lengths denote the number of nucleotide substitutions per position. Branches corresponding to non-photosynthetic species are indicated in red, and those corresponding to photosynthetic species are indicated in green. Branches in which a transition from a photosynthetic to a non-photosynthetic lifestyle occurred are indicated half in green and half in red. *Orchis italica* and *Camellia sinensis*, which are employed as outgroups in (a) and (b), respectively, are not shown, since the mutation accumulation rate of an outgroup cannot be evaluated.

### Composition of the plastid ribosome

The proteins of the plastid ribosome are encoded in both the plastid and nuclear genomes. For example, in *Arabidopsis thaliana*, 21 plastid ribosomal proteins are encoded by the plastome, and 36 are encoded by the nuclear genome. In all holo-heterotrophic plants with highly reduced plastomes, some ribosomal genes are missing; *Pilostyles aethiopica*, in which only two ribosomal protein genes are retained, represents the most extreme known case (Bellot & Renner, 2015). This raises the question of how ribosomes are able to function in these species. There are three possibilities. First, some proteins of the plastid ribosome may simply be non-essential, and their loss may not severely affect ribosome function (Tiller & Bock, 2014), although this does not explain such extreme cases of reduction. Alternatively, since the transfer and integration of plastid DNA into the nucleus exists in plant cells, functional copies of plastid genes can arise in the nuclear genome. There are examples in which the products of such nuclear copies are targeted to plastids and function as a part of the plastid ribosome, while the corresponding gene having been lost from the plastome (Ueda *et al.*, 2007; Jansen *et al.*, 2011; Park *et al.*, 2015). Additionally, components of mitochondrial ribosomes can be dually targeted to both plastids and mitochondria (Kubo & Arimura, 2010; Ueda *et al.*, 2008). In *E.aphyllum* (Schelkunov *et al.*, 2015), 7 ribosomal protein genes (*rpl20*, *rpl22*, *rpl23*, *rpl32*, *rpl33*, *rps15*, *rps16*) have been lost from the plastome; in *E.roseum* 6 of these 7 genes, but not *rpl20*, have been lost. Regarding *H.monotropa*, we previously considered *rps19* and *rpl22* to be pseudogenes (Logacheva *et al.*, 2016) due to the presence of a 111-bp insertion in the former and a nonsense mutation that shortens the length of the product by 17% in the latter. However, because these genes are transcribed and exhibit dN/dS values close to those of the species’ photosynthetic relatives (Table S7), we assume that they are functional genes. Two genes, *rps15* and *rps16*, were completely absent from the plastome of *H.monotropa*. We do not observe any transcripts with high similarity to these plastid genes in the transcriptomes of *E.aphyllum*, *E.roseum* or *H.monotropa*. Thus, the loss of these genes from the plastome is unlikely to be compensated by transfer of plastid sequences to the nuclear genome.

Instead, we found that several transcripts that are not 1-1 orthologs of plastid-encoded ones but have more distant homology to them encode proteins that can be imported into plastids. This is the case for Rpl23 and Rps15 in *E.aphyllum* and Rps15 in *H.monotropa*. Additionally, for several proteins, the predictions made with TargetP and DualPred, two tools that we used for target prediction, were contradictory. Specifically, for a homolog of Rpl32 in *E.aphyllum* and homologs of Rpl23, Rps15 and Rps16 in *E.roseum*, plastid targeting was predicted by only one of the two tools. In all cases except for Rpl23 of *E.aphyllum*, analysis of the transit peptides of the homologs in the photosynthetic relatives of these species suggested that these proteins were already targeted to plastids prior to the divergence of non-photosynthetic and photosynthetic species, which may have facilitated the loss of the respective plastid genes. Some of the aforementioned proteins are predicted to be targeted solely to plastids, and some are predicted to be dually targeted to plastids and mitochondria (for details see Table S8).

To determine whether the genes whose products may serve to replace the lost ribosomal proteins are encoded in mitochondrial or nuclear genomes, we used TBLASTN to align the proteins against contigs produced during the assembly of the plastomes of *E.aphyllum* and *H.monotropa* in our earlier studies (Schelkunov *et al.*, 2015, Logacheva *et al.* 2016, respectively). We did not observe the sequences of those genes in the mitochondrial contigs and therefore conclude that they are located in the nuclear genomes.

## Conclusions

In this study, we analysed and compared the transcriptomes of the mycoheterotrophic plants *Epipogium aphyllum, E.roseum* and *Hypopitys monotropa*. Despite the fact that *Epipogium* and *H.monotropa* are very distantly related, belonging to the Monocots and Dicots respectively, and that these species lost photosynthesis independently, we observed a remarkable level of parallelism involving the reduction and retention of similar functional groups of genes. Among the observed differences were a more profound reduction in the chlorophyll *a* synthesis pathway in *H.monotropa* and an increased rate of mutation accumulation in *Epipogium*. Overall, since there are several hundred holo-heterotrophic species of flowering plants (Merckx *et al.*, 2009), with many cases of independent transitions to holo-heterotrophic lifestyle, it is necessary to sequence and analyse more holo-heterotrophic species in addition to *Epipogium*, *H.monotropa* and *Orobanche aegyptiaca* to predict whether this parallelism is universal. Significant help in this determination may be provided by the “1000 Plants Project” (Matasci *et al.*, 2014), in which the sequencing of many holo- and hemi-heterotrophic species is being performed. The question that remains unanswered in this study is the mode of gene loss – did it occur through deletions of large regions carrying photosynthesis-related genes, or through the accumulation of mutations in the protein-coding and regulatory elements of these genes? We expect that characterization of the nuclear genomes of non-photosynthetic plants will fill this gap.

## Acknowledgements

The authors would like to thank Viktoria Shtratnikova for helpful discussion.

## Authors’ contributions

MIS performed the computational analysis and drafted the manuscript. AAP participated in sequencing. MDL sequenced the samples and participated in discussion and manuscript writing.

## Online supporting material

**Fig. S1** Statistics regarding contamination in the studied transcriptomes. A total of 10,000 random transcripts (prior to the removal of low-coverage transcripts and searching for ORFs, but after the removal of minor isoforms) were taken from each assembly, and BLASTX alignment to NCBI NR (maximum allowed e-value of 10^−5^, word size of 3 amino acids, low-complexity sequence filter switched off) was performed. The transcripts were classified according to their best matches. In the distribution plots, the black lines denote median values, and the boxes denote interquartile ranges.

**Fig. S2** Diagram of circadian rhythms regulation.

**Fig. S3** Diagram of carotenoid biosynthesis.

**Fig. S4** Diagram of ubiquinone and other terpenoid-quinone biosynthesis.

**Fig. S5** Diagram of thiamine metabolism.

**Fig. S6** Trees of the studied species with branch lengths representing dS (rate of synonymous substitutions).

**Fig. S7** Trees of the studied species with branch lengths representing dN (rate of non-synonymous substitutions).

**Fig. S8** Trees of the studied species with branch lengths representing dN/dS.

**Table S1** Information on transcriptome data.

**Table S2** Sources of genome sequences.

**Table S3** Complete list of GO terms for which the fractions of genes differed significantly among *Epipogium*, *Hypopitys monotropa* and their photosynthetic relatives * - “u” indicates underrepresentation; “o” indicates overrepresentation. Genes with a GO term are considered underrepresented in holo-heterotrophic species if their proportions relative to the total numbers of genes with GO terms in those species are significantly lower than in photosynthetic species. Overrepresentation corresponds to the opposite situation.

**Table S4** Detailed statistics of the transcriptome assemblies. * - In a dataset deposited by the authors of the *Vaccinium macrocarpon* genome assembly, several isoforms are provided for some genes. In these cases, we referred to the longest isoform as the “major” isoform, since information on the relative expression of isoforms was not supplied.

**Table S5** Analysis of the completeness of the assemblies.

**Table S6** Proportions of genes whose products are targeted to various organelles relative to the total numbers of genes in the species according to TargetP analysis. Only genes with complete 5’ ends and at least one assigned GO term are considered. The quality of the different assemblies differs; thus, the number of genes with completely assembled 5’ ends also differs. Therefore, the numbers of proteins with transit peptides cannot be directly compared, and only proportions are provided in the table.

**Table S7** Presence of genes of interest in the studied species and selective pressures acting on them.

**Table S8** Results of a search for possible substitutes for ribosomal proteins whose genes have been lost from the plastomes of *Epipogium* and *Hypopitys monotropa*.

